# Direct comparison of epifluorescence and immunostaining for assessing viral mediated gene expression in the primate brain

**DOI:** 10.1101/2022.10.04.510807

**Authors:** Tierney B. Daw, Hala G. El-Nahal, Michele A. Basso, Elizabeth J. Jun, Alex R. Bautista, R. Jude Samulski, Marc A. Sommer, Martin O. Bohlen

**Affiliations:** Department of Neurobiology, Duke University School of Medicine; Department of Biomedical Engineering, Duke University; Fuster Laboratory of Cognitive Neuroscience, Department of Psychiatry and Biobehavioral Sciences, Jane and Terry Semel Institute for Neuroscience and Human Behavior, University of California Los Angeles; Washington National Primate Research Center, Department of Biological Structure, Department of Physiology and Biophysics, University of Washington, Seattle; UNC Gene Therapy Center, University of North Carolina at Chapel Hill; Asklepios Biopharmaceutical Inc; Department of Psychology & Neuroscience, Duke University; Center for Cognitive Neuroscience, Duke University

**Author notes:** Address Correspondence to: Martin O. Bohlen, 1427 CIEMAS Building, Box 90281, 101 Science Drive, Durham, NC 27708.

**Keywords:** viral vector, non-human primate, immunofluorescence, immunohistochemistry, neuronal actuator, transgene

## Abstract

Viral vector technologies are commonly used in neuroscience research to understand and manipulate neural circuits, but successful applications of these technologies in non-human primate models have been inconsistent. An essential component to improve these technologies is an impartial and accurate assessment of the effectiveness of different viral constructs in the primate brain. We tested a diverse array of viral vectors delivered to the brain and extraocular muscles of macaques and compared three methods for histological assessment of viral-mediated fluorescent transgene expression: epifluorescence (Epi), immunofluorescence (IF), and immunohistochemistry (IHC). Importantly, IF and IHC identified a greater number of transduced neurons compared to Epi. Furthermore, IF and IHC reliably provided enhanced visualization of transgene in most cellular compartments (i.e., dendritic, axonal, and terminal fields), whereas the degree of labeling provided by Epi was inconsistent and predominantly restricted to somas and apical dendrites. Because Epi signals are unamplified (in contrast to IF and IHC), Epi likely provides a more veridical assessment for the amount of accumulated transgene and, thus, the potential to chemo- or optogenetically manipulate neuronal activity. The comparatively weak Epi signals suggest that the current generations of viral constructs, regardless of delivered transgene, are not optimized for primates. This reinforces an emerging viewpoint that viral vectors tailored for the primate brain are necessary for basic research and human gene therapy.

## INTRODUCTION

Viral vector technologies are commonly used in neuroscience research to understand and manipulate neural circuits. For example, virally delivered neuronal actuators such as opsins provide a useful way to reversibly study the role of individual neurons and circuits in the context of normal behaviors and brain disorders. Similarly, indicators such as GCaMP allow *in vivo* observation of large populations of neurons in awake, behaving animals. It is particularly valuable to use these technologies in macaques, the quintessential animal model for translational research. In macaques and other large animals, viral vectors are the standard and most effective approach for delivering exogenous genes encoding neuronal actuators and indicators. The long gestation periods and small litter sizes of macaques, along with cost and ethical considerations, hinder the development of diverse transgenic lines that are available for rats, mice, and other small lab animals. Hence, for continued progress in genetic approaches to study the primate brain and treat disease, viral delivery is central and will likely remain so for the foreseeable future.

Which viral vector works best to target a neuronal population or circuit in primates? This remains a critical question for primate brain research and has clear clinical implications for neurological gene therapies. Viral vector efficacy can be assessed in three ways: anatomical (assessing the strength and patterns of transduction via histology), neurophysiological (determining the effect on neuronal activity), and behavioral (determining the functional outcome of viral-mediated gene expression) ^1–4^. To date, all three approaches suggest that, in general, viral vector efficacy in the primate brain is relatively weak and unreliable ^2,4^. To overcome these limitations, we need to learn more about the subtleties of different viral vectors following transduction in the primate brain and determine which vector and gene constructs are most effective for driving transgene expression. Ultimately, a high level of transgene expression may be necessary to utilize optogenetic and chemogenetic tools for the manipulation of neuronal function and animal behavior. A first step toward achieving this goal is a reliable assessment of viral methods at the anatomical level to screen and benchmark a range of viral constructs that result in robust transgene expression in the primate brain.

Largely absent in the literature are comprehensive studies that compare the efficacies of multiple virus/promoter/transgene combinations to determine which, if any, have the greatest promise for use in the primate brain. Of those studies that have compared patterns of transduction, it has been commonplace to perform immunofluorescence (IF) or immunohistochemistry (IHC) despite the fact that the injected constructs contain genes encoding fluorescent reporter proteins that could be assessed directly by epifluorescence microscopy (Epi) ^5–8^. A fundamental question arises: what exactly is gained or lost by using immunostaining (IF or IHC) versus Epi to assess transduction in the primate brain? Immunostaining detects gene expression that has been amplified by antibody approaches, making it highly sensitive. One possibility is that immunostaining is more informative than Epi in every way. For example, immunostaining may reveal every dimension of transduction, including detection of gene expression in a brain area and its projections, detailed visualization of the gene product throughout cellular compartments, and predictive power about functionality of the transgene. This would suggest that, for *in vivo* experiments or gene therapies, Epi is unnecessary and anatomical reporter genes could be omitted from recombinant viral genomes that are often tight on space already (~4.7 kb for adeno-associated viruses (AAVs)). On the other hand, Epi may provide a dimension of evaluation that immunostaining cannot offer. Because it provides a raw, unamplified signal, continuing to include reporter genes in vectors may be useful for evaluating the efficacy of virally delivered therapeutic genes. Amplification by IF or IHC clearly provides greater sensitivity to the presence of the transgene, but it could provide a false impression about the level of transduction and hence the likelihood of successfully manipulating neuronal activity.

To this end, the main goal of the present study was to directly compare IF and IHC to Epi assessment of transgene expression for a wide range of viral constructs and injection sites in the macaque brain. All analyses were performed in the same laboratory to reduce confounding variables that could arise due to differences in perfusion, tissue preparation, or histological processing. Perhaps not surprisingly, IF and IHC provided a strong signal that revealed transgene expression throughout neurons and their processes (i.e., proximal and distal dendrites, axons, and terminal fields), generating the impression of high transduction efficacy. In contrast, Epi produced much weaker signals that were nearly exclusively restricted to soma and proximal dendrites, generating the impression of much less efficient transduction. These differences persisted across tested parameters including viral construct, survival duration, titer, injected volume, and injection location. The data suggest that, while many viral vectors do infect primate neurons, they likely lead to low levels of gene expression, potentially too low to reliably manipulate neuronal activity. This suggests a need to optimize these vectors to produce higher levels of transgene expression in the primate brain.

## MATERIALS AND METHODS

### Animals

All procedures were performed in accordance with the NIH *Guide for the Care and Use of Laboratory Animals* and were approved by the Institutional Animal Care and Use Committees at Duke University and the University of California, Los Angeles. Two cynomolgus macaques (*Macaca fascicularis*) and three rhesus macaques (*Macaca mulatta*) were included in this study (**Table 1**). Codes such as “M19-03” used throughout this report indicate an individual experimental animal. Some animals were previously used in transcranial magnetic stimulation studies ^9–11^ or psychophysical and electrophysiological studies ^12–16^. All animals included in this study received additional central nervous system (CNS) or peripheral nervous system (PNS) viral injections as part of different experiments ^17,18^.

**Table 1.**
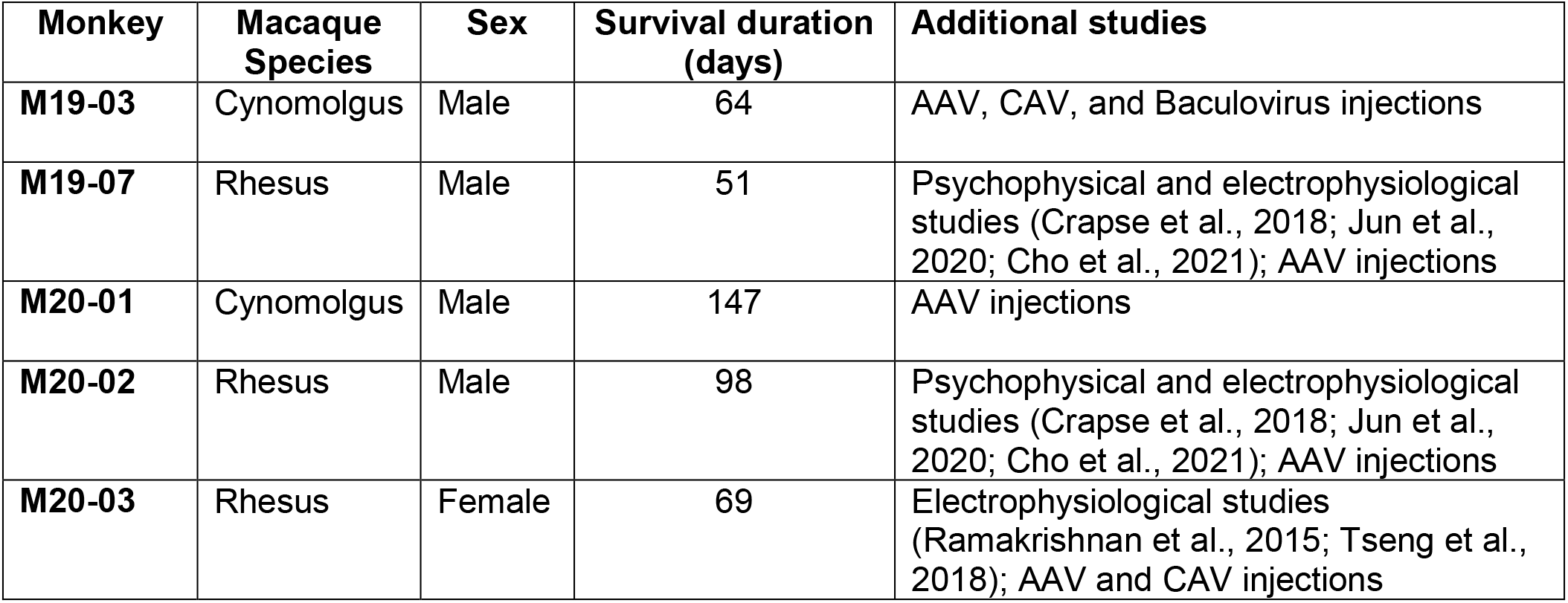
Animals included in the present study.

| Monkey | Macaque Species | Sex | Survival duration (days) | Additional studies |
| --- | --- | --- | --- | --- |
| M19-03 | Cynomolgus | Male | 64 | AAV, CAV, and Baculovirus injections |
| M19-07 | Rhesus | Male | 51 | Psychophysical and electrophysiological studies (Crapse et al., 2018; Jun et al., 2020; Cho et al., 2021); AAV injections |
| M20-01 | Cynomolgus | Male | 147 | AAV injections |
| M20-02 | Rhesus | Male | 98 | Psychophysical and electrophysiological studies (Crapse et al., 2018; Jun et al., 2020; Cho et al., 2021); AAV injections |
| M20-03 | Rhesus | Female | 69 | Electrophysiological studies (Ramakrishnan et al., 2015; Tseng et al., 2018); AAV and CAV injections |

### Viral Vectors

Viral vectors (**Table 2**) were shipped on dry ice and immediately placed in a −80°C freezer upon receipt. To avoid unnecessary freeze/thaw cycles, viral vector solutions were aliquoted following first use. Aliquots were stored in a −80°C freezer. For injection surgeries, viral solutions were stored on dry ice and thawed on wet ice prior to use, then gently centrifuged (smooth acceleration to ~3,000 RPM followed by slow deceleration) to remove any bubbles and precipitate on the sides of the centrifuge tube. Next, viral solution was gently agitated with a micropipette and drawn up into a Hamilton microsyringe or custom injectrode ^19^.

**Table 2.** Viral vectors included in the present study.

| Monkey | Viral Vector | Source | Titer (vg/ml) |
| --- | --- | --- | --- |
| M19-03 | AAV6-hSyn-hChR2(H134R)-EYFP | Duke Viral Core | $7.4 \times 10^{12}$ |
| | rAAV2-retro-hSyn-ChR2(H134R)-GFP | AddGene | $1.1 \times 10^{13}$ |
| M19-07 | rAAV2-retro-CAG-GFP | UNC Viral Core | $1.8 \times 10^{12}$ |
| M20-01 | AAV2G9-CMV-GFP (UNC) | UNC Viral Core | $3.7 \times 10^{12}$ |
| | AAV2G9-CMV-GFP (AskBio) | AskBio | $3.7 \times 10^{12}$ |
| | rAAV2-retro-CAG-tdTomato | UNC Viral Core | $2.6 \times 10^{12}$ |
| | AAV2G9<sub>(R585, 588A)</sub>-CMV-GFP | UNC Viral Core | $5.3 \times 10^{12}$ |
| M20-02 | rAAV2-retro-CAG-tdTomato | UNC Viral Core | $3.8 \times 10^{12}$ |
| M20-03 | AAV6-hSyn-hChR2(H134R)-EYFP | Duke Viral Core | $7.4 \times 10^{12}$ |

### Surgical Procedures

One day prior to surgery and on surgery day, animals were administered a corticosteroid, either Dexamethasone (2.0 mg/kg, IM) or Solu-Medrol (15.0 mg/kg, IM), for mild immunosuppression. On surgery day, animals were sedated with ketamine hydrochloride (3.0 mg/ kg, IM) and dexdomitor (0.075 mg/kg, IM). Animals were intubated and gas anesthesia was maintained throughout surgery using a 1-3% isoflurane/oxygen mix. Vital signs were monitored and maintained within normal limits by an American Veterinary Medical Associate certified veterinary technician.

All surgical procedures were performed under aseptic conditions. Planned sites for surgical incisions were thoroughly cleaned using betadine, chlorohexidine, then 100% ethanol scrubs.

An injection of ~1 mL 0.25% bupivacaine was cutaneously administered along the incision line prior to all incisions and following final suturing. Animals received buprenorphine SR (0.2 mg/kg, IM) for post-operative analgesia. Dexamethasone was administered for 10 days post-operatively, starting at a dosage of 2.0 mg/kg and tapered to 0.5 mg/kg; alternatively, Solu-Medrol was administered for 10-days post-op, starting at a dosage of 15.0 mg/kg and tapered to 1.0 mg/kg.

#### Central injections

**Table 3** lists the viral injections that were included in this study. For cortical injections (M19-03 and M20-01), animals were placed into a stereotaxic apparatus (Kopf Instruments, Tujunga, CA). A midline incision was made, and soft tissues were retracted to visualize the skull. Stereotaxic coordinates were used to determine the location of the targeted brain region ^20^. A craniotomy and durotomy were performed to reveal the underlying cortex. The location of each targeted cortical region was confirmed visually. After making a surface measurement, a Hamilton syringe held in a micromanipulator was advanced to a predetermined depth below the cortical surface. Virus was then dispensed using pressure injection at a rate of 0.5 or 1 μl/min, followed by a 2- to 10-minute waiting period to allow virus to diffuse from the injection site. For some injections, virus was distributed at several sites throughout the targeted brain structure. M19-03 was injected with 25 μl of rAAV2-retro-hSyn-ChR2(H134R)-GFP in left frontal eye field (FEF). M20-01 received 4 μl of AAV2G9-CMV-GFP (UNC) in left parietal area, PE (PE)/medial intraparietal area (MIP), 5 μl of AAV2G9-CMV-GFP (AskBio) in right PE/MIP, 5 μl of rAAV2-retro-CAG-tdTomato in left FEF, and 5 μl of AAV2G9_(R585, 588A)_-CMV-GFP in right FEF. Following injections, surgical sites were thoroughly flushed with sterile saline, dura was sutured back together, and the bone flap was replaced. Soft tissues and skin were reapproximated and sutured.

**Table 3.** Viral injections included in the present study.

| Monkey | Viral Vector | Total Injection Volume | Number of injections | Rate (μl/min) | Injection location |
| --- | --- | --- | --- | --- | --- |
| M19-03 | AAV6-hSyn-hChR2(H134R)-EYFP | 100 μl | 2 (83 μl + 17 μl) | N/A | Left medial rectus |
|  | rAAV2-retro-hSyn-ChR2(H134R)-GFP | 25 μl | 10 (2.5 μl/injection) | 1 | Left FEF |
| M19-07 | rAAV2-retro-CAG-GFP | 3 μl | 1 | 0.1 | Right SC |
| M20-01 | AAV2G9-CMV-GFP (UNC) | 4 μl | 1 | 0.5 | Left PE/MIP |
|  | AAV2G9-CMV-GFP (AskBio) | 5 μl | 2 (2.5 μl/injection) | 0.5 | Right PE/MIP |
|  | rAAV2-retro-CAG-tdTomato | 5 μl | 1 | 0.5 | Left FEF |
|  | AAV2G9<sub>(R585, 588A)</sub>-CMV-GFP | 5 μl | 1 | 0.5 | Right FEF |
| M20-02 | rAAV2-retro-CAG-tdTomato | 3 μl | 1 | 0.1 | Right SC |
| M20-03 | AAV6-hSyn-hChR2(H134R)-EYFP | 80 μl | 1 | N/A | Left medial rectus |

Monkeys M19-07 and M20-02 had existing chambers, which provided access to the superior colliculus (SC). Electrophysiologically identified coordinates were used to locate the SC. A custom injectrode was used. The injectrode was inserted through a guide tube positioned inside the chamber. Virus was pressure injected at a rate of 0.1 μl/min and allowed to diffuse from the injection site for 10 min. M19-07 received 3 μl of rAAV2-retro-CAG-GFP in the right SC. M20-02 received 3 μl of rAAV2-retro-CAG-tdTomato in the right SC.

#### Peripheral injections

Extraocular muscle injections were performed as previously described (Bohlen et al., 2019). M19-03 received 100 μl of AAV6-hSyn-hChR2(H134R)-EYFP while M20-03 received 80 μl of AAV6-hSyn-hChR2(H134R)-EYFP, both in the left medial rectus.

### Histology

Following a survival duration (**Table 1**), animals were sedated with ketamine hydrochloride (3.0 mg/kg, IM), then given a lethal injection of sodium pentobarbital (50.0 mg/kg, IP). Once areflexic, animals were transcardially perfused first with 1-4 L of chilled 0.1 M, pH 7.4 phosphate buffered saline (PBS), followed by 4 L of 4% paraformaldehyde (PFA) in PBS. The brain was cut into 2-3 cm blocks *in situ* using a stereotaxic apparatus. Blocks were post-fixed in 4% PFA/PBS at 4°C for 24-48 hours, then cryoprotected in 30% sucrose in PBS at 4°C. Once blocks sank in sucrose, they were sectioned into 75 μm thick coronal sections using a freezing stage sliding microtome (American Optical Company, Buffalo, NY). Sections were stored in PBS at 4°C. For processing, sections were divided into 6 rostral to caudal series, with ~450 μm between adjacent sections within a series.

#### Processing for epifluorescence microscopy

In each case, viral-mediated Epi was observed in the first series without immunological amplification. Immediately after sectioning, the first series was mounted on 10% gelatinized glass slides and dried overnight. Mounted sections were briefly rinsed in PBS and dehydrated in a gradient of alcohols, then in xylene or toluene. Slides were coverslipped using Cytoseal 60 (Thermo Fisher Scientific, Waltham, MA).

#### Immunofluorescence processing

Serial free-floating sections were rinsed in PBS, then bathed in a Triton solution (0.5-0.75% Triton X-100 in PBS). To prevent non-specific antibody binding, sections were incubated in Bovine Serum Albumin (BSA) solution (1% BSA and 0.5-0.75% Triton X-100 in PBS). Sections were incubated on a shaker plate in antibody solutions against the appropriate reporter genes, described below.

Dual IF was performed for visualization of red (tdTomato) and green (GFP or YFP) reporter gene expression. Primary antibody solutions contained ~1:200 dilution of primary antibody, 1% BSA, and 0.5-0.75% Triton X-100 in PBS. Sections were incubated at 4°C for ~3 days in primary antibody solution containing rabbit anti-RFP (Rockland Immunochemicals Inc., Limerick, PA; 600-401-379) and biotinylated goat anti-GFP (Rockland; 600-106-215). Next, sections were washed with PBS. IF secondary antibody solutions contained ~1:200 dilution of secondary antibody, 0.5% Triton X-100, and 9% normal donkey serum in PBS. Sections were incubated at room temperature for ~3 hours in secondary antibody solution containing donkey anti-rabbit IgG H&L Texas Red (abcam, Cambridge, UK; ab7081) and donkey anti-goat IgG H&L Alexa Fluor 488 (abcam; ab150129).

Alternatively, to compare Epi to IF amplification in the same tissue section, an opposite color secondary fluorophore was used. Sections were incubated at 4°C for ~3 days in primary antibody solution containing rabbit anti-RFP (Rockland; 600-401-379). Next, sections were washed with PBS. Sections were incubated at room temperature for ~3 hours in secondary antibody solution containing goat anti-rabbit IgG H&L Alexa Fluor 488 (abcam; ab150077).

Following incubation in secondary antibody, sections were washed with PBS, then mounted, dehydrated, and coverslipped as described above.

#### Immunohistochemical processing

Serial free-floating sections were washed with PBS, then incubated in hydrogen peroxide solution (0.3-3% H_2_O_2_ in PBS) to block endogenous peroxidase activity. Following a PBS wash, sections were washed in Triton solution, then incubated in BSA solution. Sections were incubated on a shaker plate in antibody solutions against the appropriate reporter genes, described below.

Sections were incubated at 4°C for ~3 days in primary antibody solution containing either biotinylated goat anti-GFP (Rockland; 600-106-215) or peroxidase conjugated rabbit anti-RFP (Rockland; 600-403-379) for visualization of green and red fluorescent protein expression, respectively. Next, sections were washed with PBS. IHC secondary antibody solutions contained 1:200 dilution of secondary antibody and 1.5% normal rabbit or goat serum in PBS. Sections were incubated at room temperature for 1.5 hours in secondary antibody solution containing either biotinylated rabbit anti-goat IgG (Vector Laboratories, Burlingame, CA; PK-6105) or biotinylated goat anti-rabbit IgG (Vector Laboratories; PK-6101) to amplify green and red gene expression, respectively. Sections were washed in PBS, then incubated in Avidin-Biotin-horseradish peroxidase Complex (Vector Laboratories; PK-6101 and PK-6105) in PBS. Following a PBS wash, sections were transferred to diaminobenzidine tetrahydrochloride (DAB) solution (0.016% DAB, 0.016% cobalt chloride, and 0.016% nickel ammonium sulfate in Phosphate Buffer (0.4 M, pH 7.2)). After incubation in DAB solution, hydrogen peroxide solution was added to catalyze the chromogenic reaction. Sections were washed in PBS, mounted on glass slides, and dried overnight. Mounted sections were washed in PBS, counterstained using 0.04% thionin, then dehydrated and coverslipped as described above.

### Analysis

Bright field and fluorescence digital photomicrographs were taken on a Zeiss Axio Scan.Z1 using a Hitachi HV F202 color camera and Zeiss Axiocam 506 mono for fluorescent photomicrographs (Carl Zeiss Microscopy, LLC, White Plains, NY). The microscope and cameras were controlled by Zeiss ZEN 2.3 software. Within each figure, camera settings and exposure times were kept constant to avoid the introduction of any bias caused by differences in camera settings across images. When necessary, bright field and fluorescence images were adjusted for brightness, contrast, and color using Photoshop (Adobe Systems Inc., San Jose, CA) to replicate the images as they appeared when visualized under the microscope. When image adjustment was performed, all photomicrographs for a case (i.e., Epi and IF) were adjusted as a single image to prevent bias in visualizing fluorescent protein expression. Corel Draw (Corel Corp., Ottawa, Ontario) was used to reconstruct gross structural anatomy.

## RESULTS

The present study includes viral injections made in five macaques (**Table 1**). Viral vectors included several AAV serotypes, which were neuroanatomically evaluated for their propensity to transduce and drive transgene expression within the CNS. Viral vector and injection details are listed in **Tables 2 and 3**, respectively.

**Figure 1** illustrates example CNS injection sites. Photomicrographs show the locations of injections in the coronal plane and the degree of viral mediated labeling following Epi assessment. The Epi signal was frequently faint. Injections of rAAV2-retro-hSyn-ChR2(H134R)-GFP were placed in the left FEF of M19-03 (**Fig. 1a**). These injections were concentrated medially within the FEF with identifiable, but weak labeling spanning cortical layers. Injections of rAAV2-retro-CAG-GFP (M19-07) (**Fig. 1b**) and rAAV2-retro-CAG-tdTomato (M20-02) (**Fig. 1c**) were placed in the right SC. In both cases, SC labeling was most concentrated within the intermediate layers with more diffuse labeling extending ventrally into the deep and dorsally into the superficial tectal layers. In M20-01, rAAV2-retro-CAG-tdTomato was injected in the left FEF filling all lamina (**Fig. 1d**). In the same animal and during the same surgery, AAV2G9_(R585, 588A)_-CMV-GFP was placed in the right dorsal portion of FEF (**Fig. 1e**) and AAV2G9-CMV-GFP was injected into the left and right PE/MIP (**Fig. 1f**), but these injections yielded scant labeling across cortical layers. Also included in the study were peripheral muscle injections of AAV6-hSyn-hChR2(H134R)-EYFP in the left medial rectus (M19-03 and M20-03) (injection sites not shown).

**Figure 1.**
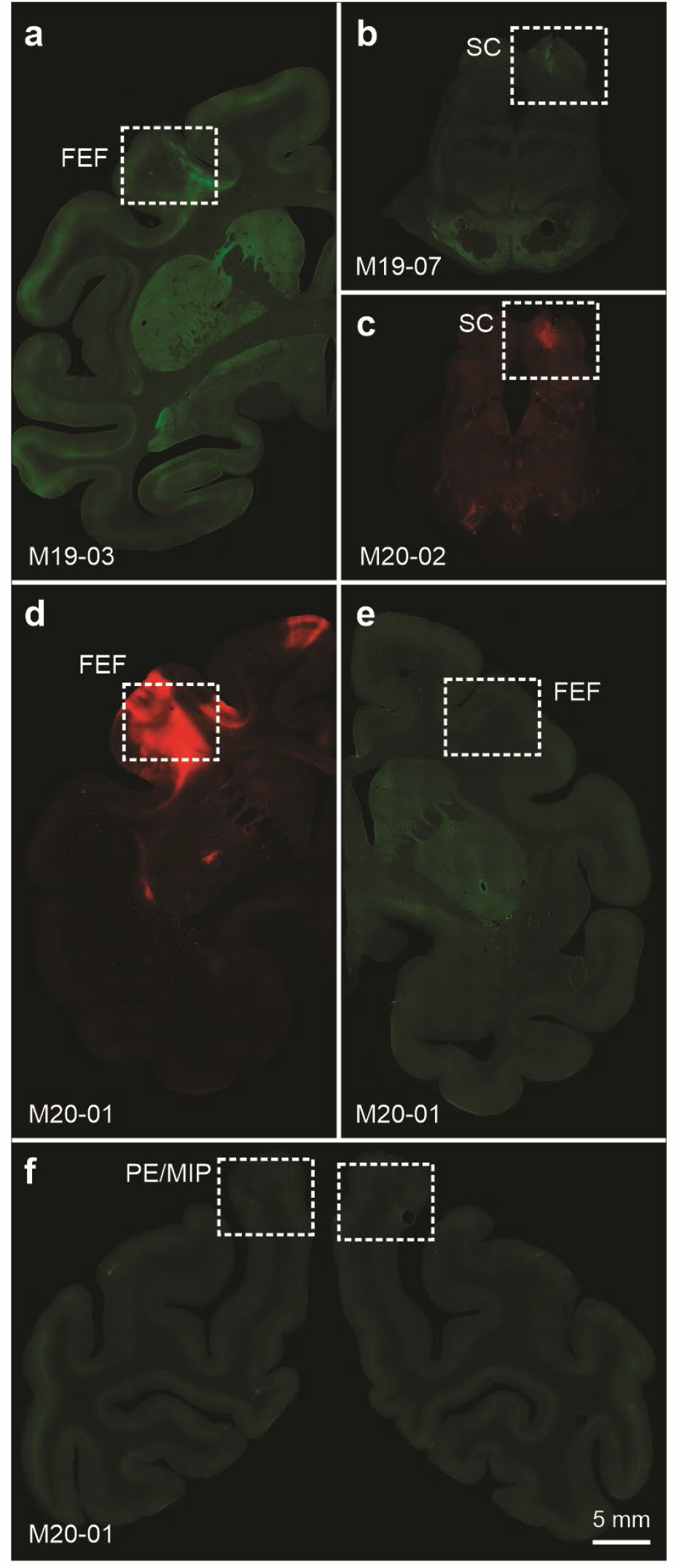
Epi visualization of CNS injection sites. (**a**) Injections of rAAV2-retro-hSyn-ChR2(H134R)-GFP in left FEF (M19-03). (**b**) Injections of rAAV2-retro-CAG-GFP (M19-07) and (**c**) rAAV2-retro-CAG-tdTomato (M20-02) in right SC. (**d**) Injection of rAAV2-retro-CAG-tdTomato in left FEF, (**e**) AAV2G9_(R585, 588A)_-CMV-GFP in right FEF, and (**f**) AAV2G9-CMV-GFP in left and right PE/MIP (M20-01). The scale bar in 1f applies to all photomicrographs.

### Anatomical assessment of local transduction and expression at the injection site

First, different AAV viral constructs were evaluated for their ability to drive transgene expression at the injection sites by comparing Epi to immunostained tissue. **Figures 2 and 3** provide low magnification drawings illustrating injection site placement and photomicrographs that show labeling visualized using Epi, IF, and IHC for intraparenchymal injection sites. Epi assessments of transduction and fluorescent reporter protein expression were highly variable across virus-type, promoter, and fluorescent protein. One injection used rAAV2-retro-hSyn-ChR2(H134R)-GFP in the left FEF (**Fig. 2a-g**). Epi (**Fig. 2b, e**) showed only moderate labeling of proximal dendrites (**Fig. 2e; black arrowheads**) of pyramidal neurons local to the injection site. However, when neighboring sections were analyzed with IF (**Fig. 2c, f**) or IHC (**Fig. 2d, g**), the amplification produced a robust signal in proximal dendrites (**Fig. 2f, g; black arrowheads**) and provided visualization of the distal stem and dendritic tufts (**Fig. 2f, g; white arrowheads**) in the superficial cortical layers.

**Figure 2.**
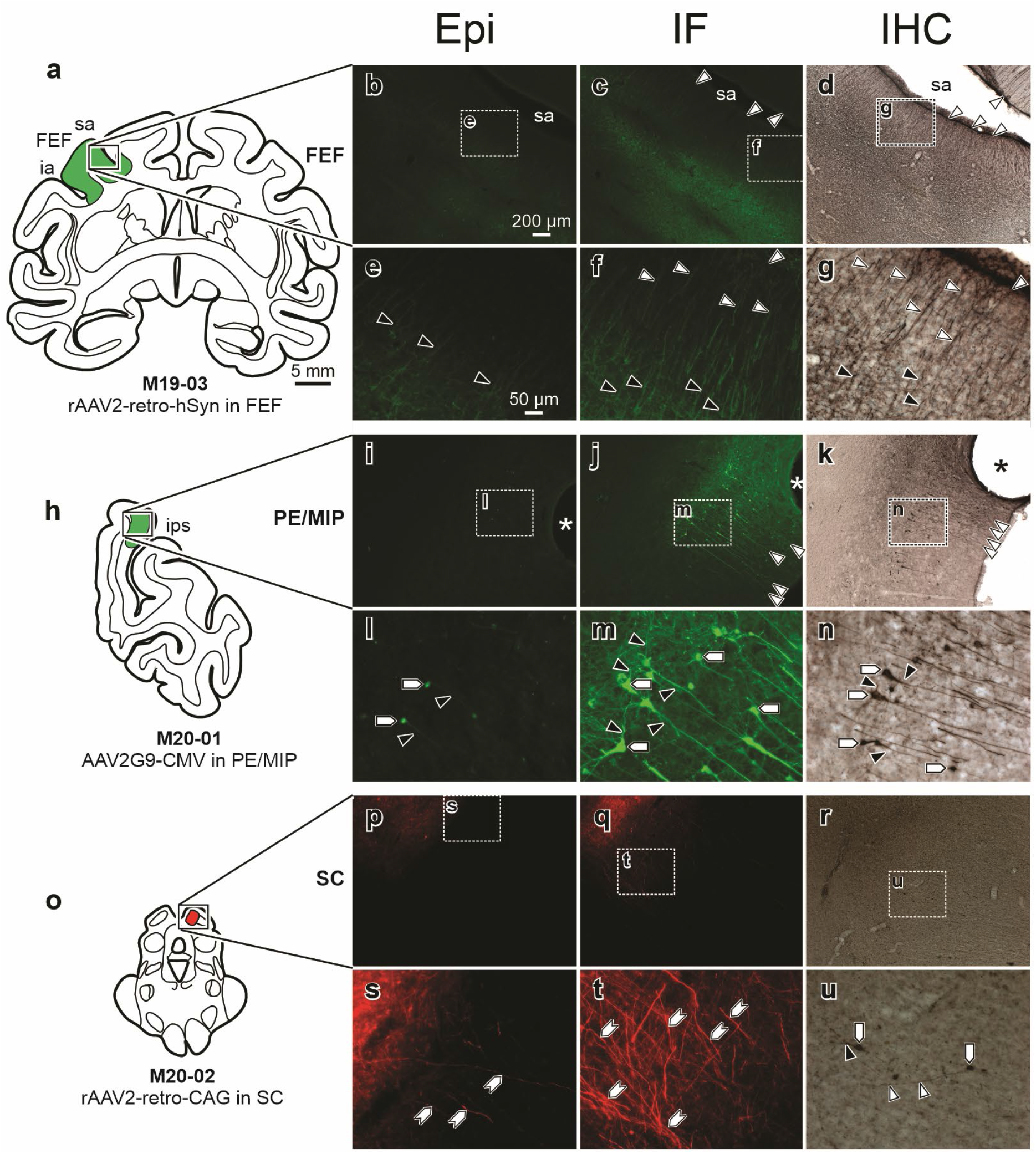
Moderate fluorescent reporter transgene expression was observed in Epi sections compared with robust detectable transgene expression following IF and IHC amplification at central injection sites. (**a**) Low magnification drawing illustrating the injection of rAAV2-retro-hSyn-ChR2(H134R)-GFP in left FEF of M19-03. Photomicrographs from neighboring sections of the left FEF are shown to visualize Epi (**b, e**) or tissue reacted for IF (**c, f**) or IHC (**d, g**). (**h**) Low magnification drawing of the AAV2G9-CMV-GFP injection in right PE/MIP in M20-01. Photomicrographs include Epi (**i, l**), IF (**j, m**), and IHC (**k, n**) imaged from neighboring sections. For subcortical representation, (**o**) illustrates a low magnification drawing of the rAAV2-retro-CAG-tdTomato injection in right SC of M20-02. Photomicrographs include Epi (**p, s**), IF (**q, t**), and IHC (**r, u**) imaged from neighboring sections. Labeled soma are indicated with **pentagons**, proximal dendrites are indicated by **black arrowheads**, and distal stem and dendritic tufts by **white arrowheads**. Since it is unclear what portion of the neuronal cytoarchitecture is labeled in the SC case, **hexagons** are used to indicate general labeling of neuronal processes. Scale bar in **a** (5 mm) applies to **h** and **o**; scale bar in **b** (200 μm with 5× objective) applies to **c-d**, **i-k**, and **p-r**; scale bar in **e** (50 μm at 20×) applies to **f-g**, **l-n**, and **s-u**. GFP and RFP were visualized with fluorescence microscopy for Epi and IF or bright field microscopy for IHC. Z-stacks: e = 15z; f = 15z; g = 18z; l = 15z; m = 13z; n = 13z; s = 16z; t = 18z; u = 20z.

**Figure 3.**
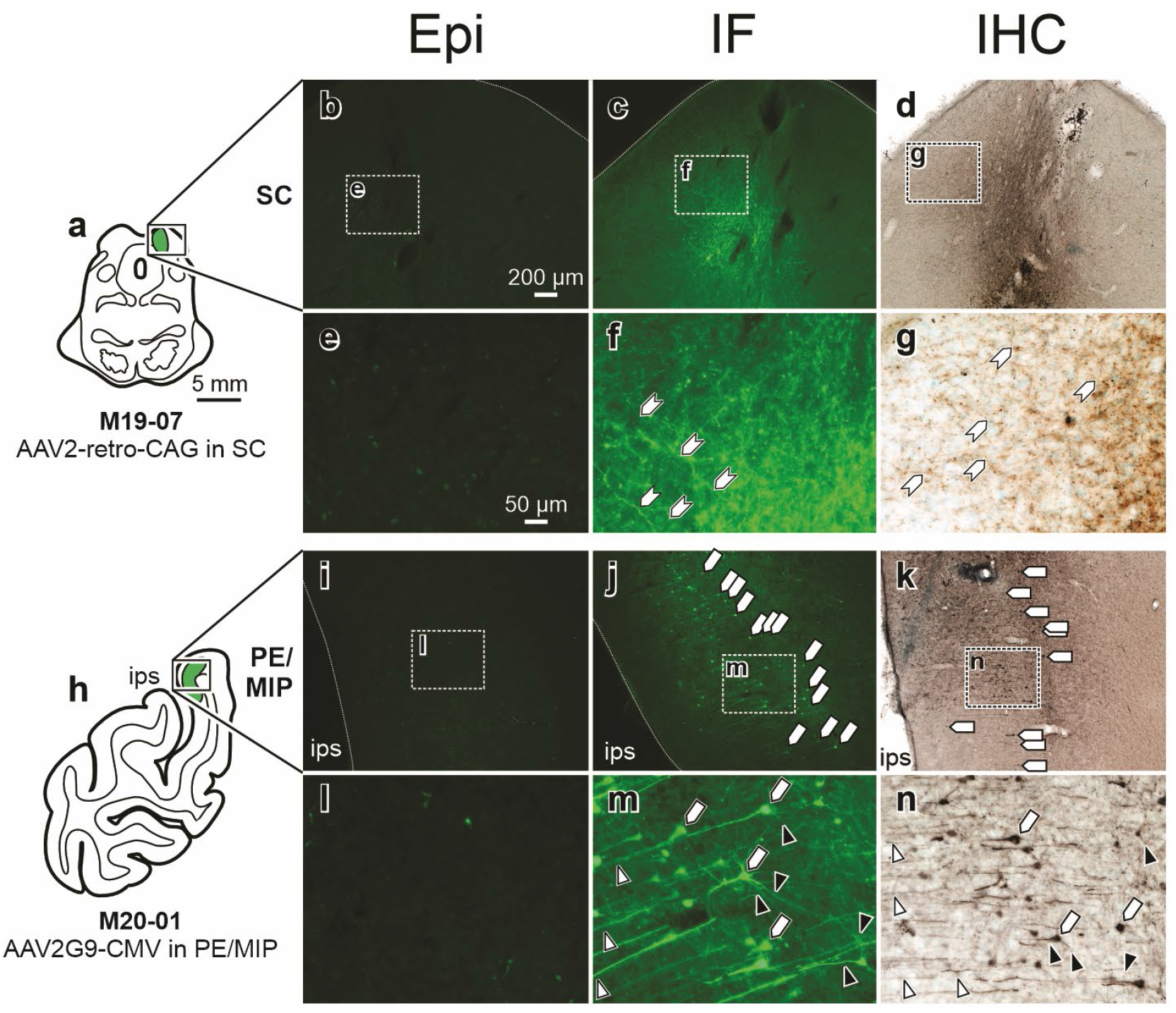
Minimal GFP labeling was visualized with Epi, whereas IF and IHC revealed robust transgene expression at central injection sites. (**a**) Low magnification drawing of rAAV2-retro-CAG-GFP injection in right SC of M19-07. Epi (**b, e**), IF (**c, f**), and IHC (**d, g**) sections for M19-07 SC. Only IF and IHC provided visualization of labeling in small neuronal processes (**hexagons**). (**h**) Low magnification drawing of AAV2G9-CMV-GFP injection in left PE/MIP of M20-01. Epi (**i, l**), IF (**j, m**), and IHC (**k, n**) for M20-01 PE/MIP. IF and IHC revealed profuse labeling in soma (**pentagons**), proximal (**black arrowheads**), and distal dendrites (**white arrowheads**). Scale bar in **a** (5 mm) applies to **h**; scale bar in **b** (200 μm at 5×) applies to **c-d** and **i-k**; scale bar in **e** (50 μm at 20×) applies to **f-g** and **l-n**. GFP was visualized with fluorescence microscopy for Epi and IF or bright field microscopy for IHC. Z-stacks: e = 16z; f = 24z; g = 28z; l = 16z; m = 14z; n = 15z.

Similarly, an injection of AAV2G9-CMV-GFP in right PE/MIP (**Fig. 2h-n**) resulted in Epi detection (**Fig. 2i, l**) of few pyramidal neuron somas (**Fig. 2l; pentagons**) and proximal dendrites (**Fig. 2l; black arrowheads**) at the injection site. Again, IF (**Fig. 2j, m**) and IHC (**Fig. 2k, n**) in neighboring sections produced greater detail in terms of the morphology of individual neurons, providing visualization of viral expression in somas (**Fig. 2m, n; pentagons**), basal and proximal dendrites (**Fig. 2m, n; black arrowheads**), and the distal stem and dendritic tufts (**Fig. 2j, k; white arrowheads**).

To illustrate that this phenomenon is not exclusive to the cerebral cortex, a third case is presented where the right SC (**Fig. 2o-u**) was injected with rAAV2-retro-CAG-tdTomato. This injection resulted in moderate Epi labeling (**Fig. 2p, s**) of neuronal processes (**Fig. 2s; hexagons**). In comparison, IF (**Fig. 2q, t**) produced dense labeling of neuronal processes (**Fig. 2t; hexagons**) at the injection site, and IHC (**Fig. 2r, u**) produced labeling in somas (**Fig. 2u; pentagons**), proximal dendritic fields (**Fig. 2u; black arrowheads**), and distal dendritic fields (**Fig. 2u; white arrowheads**). Overall, compared to Epi assessment, immunostaining provided particularly excellent visualization of small neuronal processes and a greater number of labeled cell bodies.

Of particular concern were cases exhibiting minimal detectable fluorescent labeling at the injection site using Epi. Following an injection of rAAV2-retro-CAG-GFP in the right SC (**Fig. 3a-g**), negligible labeling was visualized at the injection site using Epi (**Fig. 3b, e**), while IF (**Fig. c, f**) and IHC (**Fig. d, g**) amplified the signal to show robust viral mediated GFP expression. Specifically, small neuronal processes (**Fig. 3f, g; hexagons**) were detectable at the SC injection site using IF and IHC, but not Epi. In another case, when AAV2G9-CMV-GFP was injected in the left PE/MIP (**Fig. 3h-n**), little Epi labeling was observed at the injection site (**Fig. 3i, l**). Following IF (**Fig. 3j, m**) or IHC (**Fig. 2k, n**), somas (**Fig. 3j, k, m, n; pentagons**), basal and proximal dendrites (**Fig. 3m, n; black arrowheads**), and distal dendritic fields (**Fig. 3m, n; white arrowheads**) were visible at the injection site in PE/MIP.

Together, Figures 2 and 3 demonstrate that Epi assessment was generally insufficient for determining patterns of transduction and transgene expression at the injection site, whereas IF and IHC permitted visualization of viral mediated expression in most parts of the neuron, especially small neuronal processes such as dendritic fields.

### Anatomical assessment of transgene expression in anterogradely filled axons and axon terminals arising from neurons transduced at the injection site

Although directly evolved in rodents for retrograde axonal transport, rAAV2-retro also provides anterograde axonal fill in macaques and may be useful for studying efferent projections ^18^. While the mediodorsal thalamic nucleus (MD) is reciprocally connected to FEF, rAAV2-retro fails to retrogradely transduce MD thalamocortical neurons following injections in FEF ^17,18^. In contrast, MD terminal field labeling was observed, putatively from corticothalamic neurons that have been transduced in FEF ^18^. Following an injection of rAAV2-retro-hSyn into the left FEF (**Fig. 4a**), we looked at two major recipients of FEF input, MD (**Fig. 4b-h**) and SC (**Fig. 4i-o**). Despite being a well-established corticothalamic projection, injections of rAAV2-retro-hSyn-ChR2(H134R)-GFP in left FEF revealed minimal terminal field labeling within left MD (**Fig. 4b**) upon Epi assessment (**Fig. 4c, f**). However, IF (**Fig. 4d, g**) and IHC (**Fig. 4e, h**) both revealed densely labeled axonal terminal fields (**Fig. 4g, h; arrows**) in MD in adjacent sections. From the same case, we also examined the left SC (**Fig. 4i**) for evidence of FEF terminal field labeling. Labeling was largely absent using Epi (**Fig. 4j, m**), but transgene expression in terminal fields (**Fig. 4n, o; arrows**) was evident using IF (**Fig. 4k, n**) and IHC (**Fig. 4l, o**). To determine if this was a function of the rAAV2-retro or a general feature of AAVs, we examined the SC (**Fig. 4q**) for terminal field labeling following injections of rAAV2G9-CMV-GFP into the FEF (**Fig. 4p**). Similar to what was observed with rAAV2-retro, Epi assessment of rAAV2G9 revealed negligible axon terminal labeling in the intermediate and deep tectal layers (**Fig. 4r, u**). However, IF (**Fig. 4s, v**) and IHC (**Fig. 4t, w**) showed profuse labeling in terminal fields (**Fig. 4v, w; arrows**).

**Figure 4.**
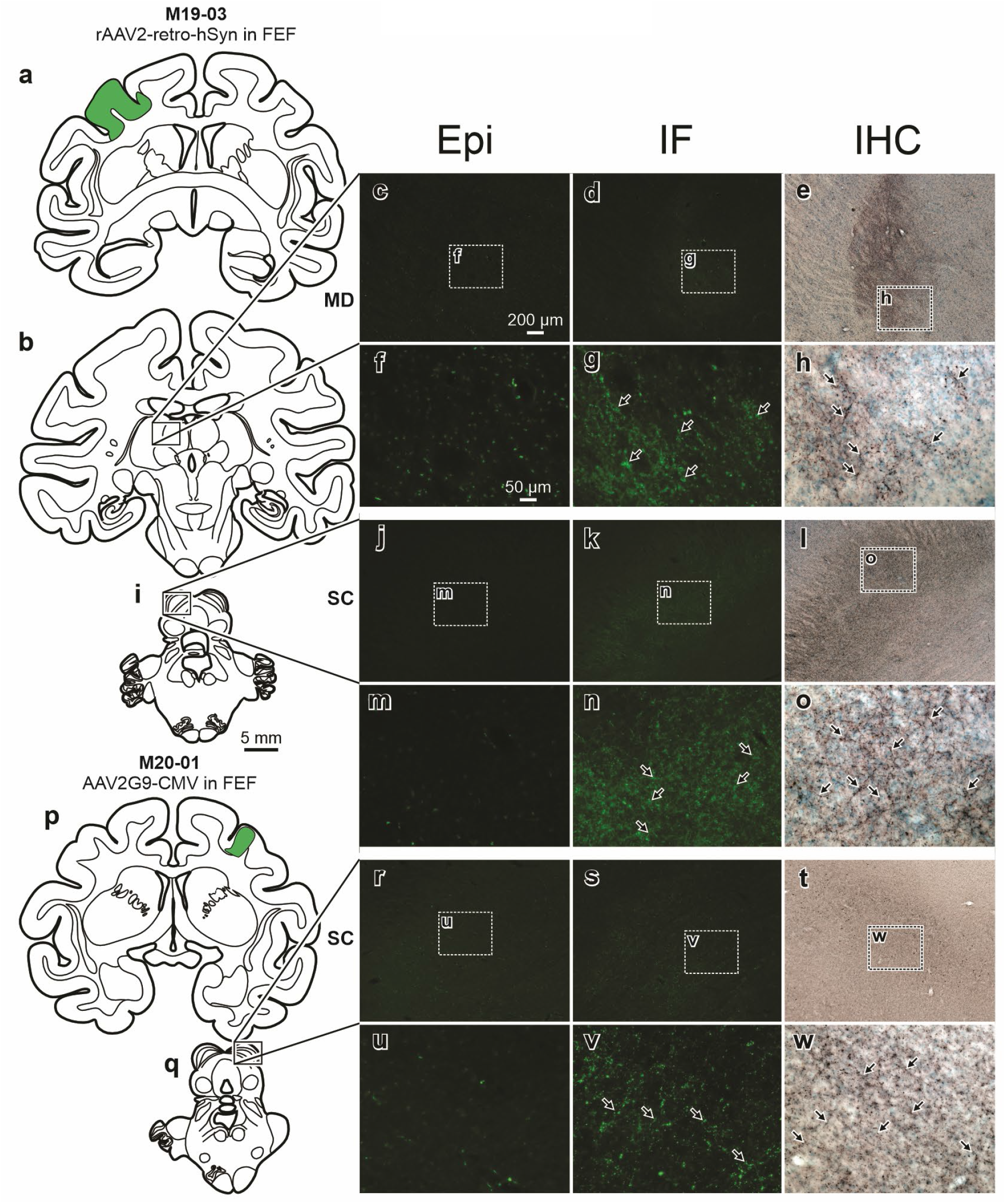
Anterograde fill: transgene expression was minimally detectable in Epi, but robust in IF and IHC. (**a**) Low magnification drawings illustrate the rAAV2-retro-hSyn-ChR2(H134R)-GFP injection site in the left FEF. This injection resulted in terminal field labeling in (**b**) MD and (**i**) SC. (**b**) Low magnification drawing illustrating the approximate location of the left MD photomicrographs (**c-h**) from case M19-03. Epi (**c, f**), IF (**d, g**), and IHC (**e, h**) sections for M19-03 MD. (**i**) Low magnification drawing illustrating the approximate location of the left SC photomicrographs from the same FEF injection illustrated in (**a**). Epi (**j, m**), IF (**k, n**), and IHC (**l, o**) sections for M19-03 SC. (**p**) Low magnification drawing illustrating the rAAV2G9-CMV-GFP injection site into right FEF. (**q**) Low magnification drawing illustrating the approximate location of the right SC photomicrographs. Epi (**r, u**), IF (**s, v**), and IHC (**t, w**) sections for M20-01 SC. In all cases, robust labeling of axonal terminal fields (**arrows**) was observed only when IF or IHC assessment was employed. Scale bar in **i** (5 mm) applies to **a-b** and **p-q**; scale bar in **c** (200 μm at 5×) applies to **d-e**, **j-l**, and **r-t**; scale bar in **f** (50 μm at 20×) applies to **g-h**, **m-o**, and **u-w**. GFP was visualized with fluorescence microscopy for Epi and IF or bright field microscopy for IHC. Z-stacks: e = 24z; f = 21z; g = 23z; l = 19z; m = 20z; n = 10z; s = 16z; t = 18z; u = 15z.

### Anatomical assessment of transgene expression within retrogradely transduced neuronal populations

Neuroanatomical connections between visual and visuomotor regions within the monkey CNS have been well-established using conventional tracers. The ability to retrogradely deliver and express transgenes encoding actuators or indicators within neurons is fundamental for projection targeting with phototagging experiments ^21–23^. Therefore, we next evaluated the ability for retrograde AAVs to deliver and drive fluorescent transgene expression in neurons with projections to the injection sites. In the CNS, we used rAAV2-retro, capitalizing on projections for which it is known to provide efficient retrograde transduction ^18^. In peripheral muscle injections, we used AAV6. Overall, Epi detection of retrogradely delivered fluorescent reporter protein expression ranged from moderate (**Fig. 5**) to undetectable (**Fig. 6 and 7**). For instance, following an injection of rAAV2-retro-CAG-tdTomato in the right SC (**Fig. 5a**), retrograde labeling within secondary visual cortex (V2) (**Fig. 5b-h**) was examined. Epi assessment (**Fig. 5c, f**) showed moderate retrograde labeling of soma (**Fig. 5f; pentagon**) and proximal and basal dendrites (**Fig. 5f; black arrowheads**) in V2. However, IF (**Fig. 5d, g**) and IHC (**Fig. 5e, h**) enhanced labeling of soma (**Fig. 5d, e, g, h; pentagons**) and allowed visualization of a high density of proximal dendritic fields (**Fig. 5g, h; black arrowheads**), the distal stem (**Fig. 5d, e; white arrowheads**), and dendritic tufts.

**Figure 5.**
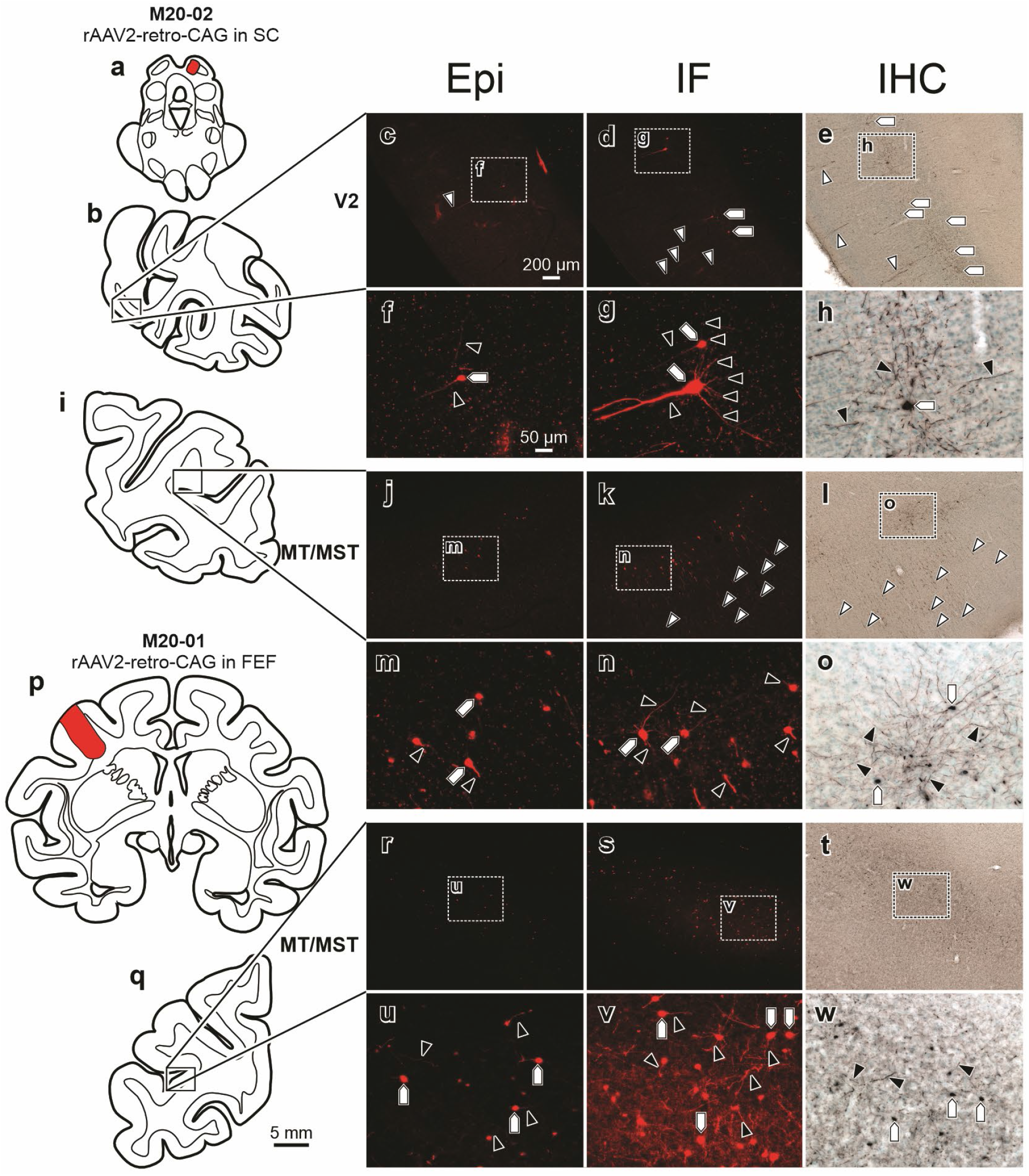
Retrograde fill: moderate RFP labeling was observed in retrogradely transduced neuronal populations in Epi sections, compared to robust labeling observed with IF and IHC. (**a**) Low magnification drawing illustrating the injection of AAV2-retro-CAG-tdTomato in right SC of M20-02. Examples of resulting retrograde labeling are illustrated within V2 (**b-h**) and MT/MST (**i-o**). (**b**) Low magnification, representative drawing illustrating the approximate location of V2 where photomicrographs were taken from neighboring sections that have been processed for Epi (**c, f**), IF (**d, g**), and IHC (**e, h**). (**i**) Low magnification, representative drawing illustrating the approximate location of MT/MST where photomicrographs were taken from neighboring sections that have been processed for Epi (**j, m**), IF (**k, n**), and IHC (**l, o**). (**p**) Low magnification drawing that illustrates the injection of AAV2-retro-CAG-tdTomato in left FEF of M20-01. (**q**) Low magnification, representative drawing illustrating the approximate location of MT/MST where photomicrographs were taken from neighboring sections that have been processed for Epi (**r, u**), IF (**s, v**), and IHC (**t, w**). In all cases, IF and IHC processing provided enhanced visualization of soma (**pentagons**) and proximal dendritic field (**black arrowheads**) labeling. Scale bar in **q** (5 mm) applies to **a-b**, **i**, and **p**; scale bar in **c** (200 μm at 5×) applies to **d-e**, **j-l**, and **r-t**; scale bar in **f** (50 μm at 20x) applies to **g-h**, **m-o**, and **u-w**. Z-stacks: e = 15z; f = 15z; g = 20z; l = 17z; m = 12z; n = 26z; s = 15z; t = 13z; u = 23z.

**Figure 6.**
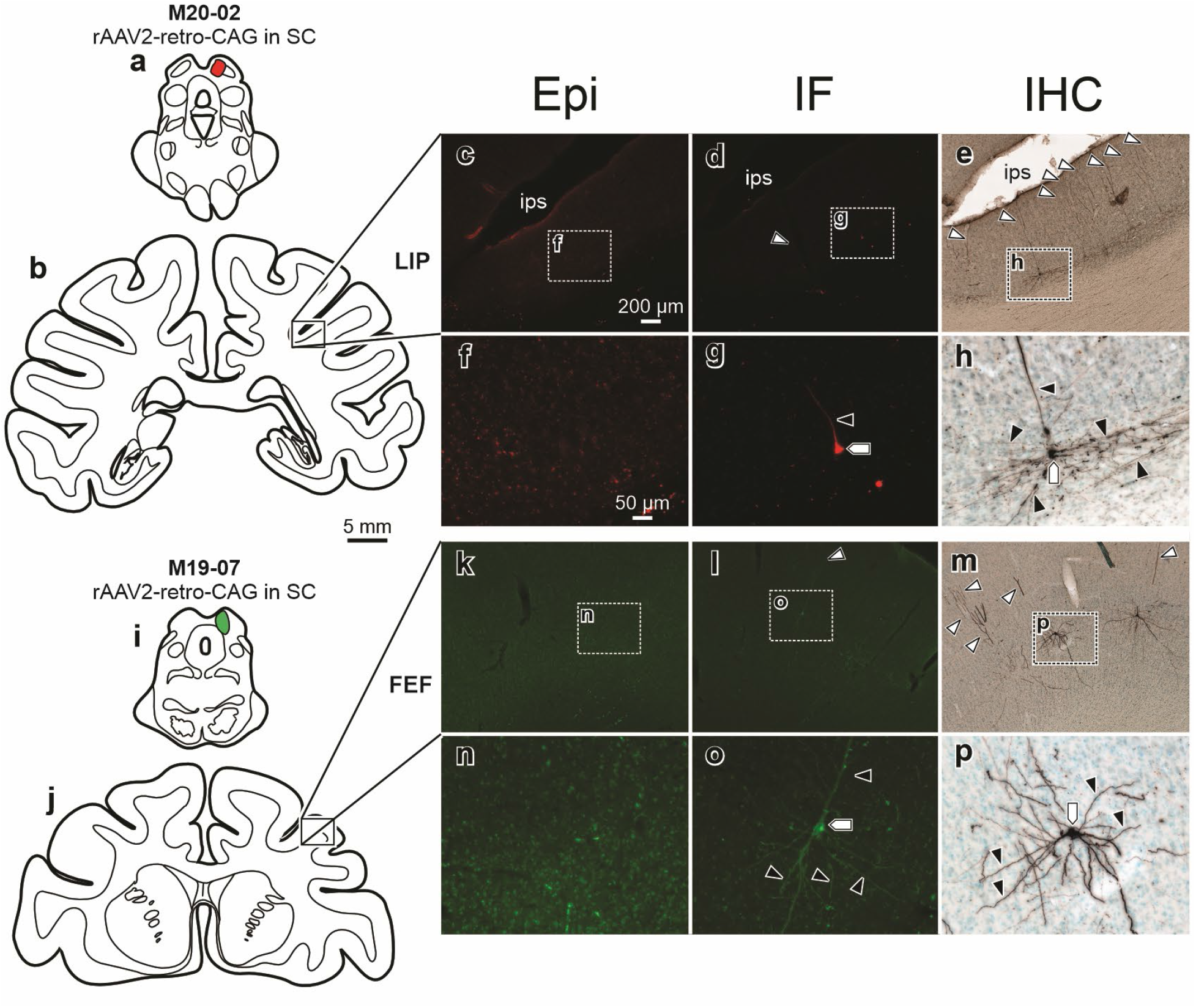
Retrograde fill: minimal transgene expression was detectable in Epi sections compared to IF and IHC, which revealed retrograde labeling at projections. (**a**) Low magnification drawing illustrating the injection of rAAV2-retro-CAG-tdTomato in right SC of M20-02. (**b**) Low magnification, representative drawing illustrating the approximate location of LIP where photomicrographs were taken from neighboring sections that have been processed for Epi (**c, f**), IF (**d, g**), and IHC (**e, h**). (**i**) Low magnification drawing illustrating the injection of rAAV2-retro-CAG-GFP in right SC of M19-07. (**j**) Low magnification, representative drawing illustrating the approximate location of FEF where photomicrographs were taken from neighboring sections that have been processed for Epi (**k, n**), IF (**l, o**), and IHC (**m, p**). In both examples, transgene expression was undetectable using Epi, whereas IF and IHC provided visualization of labeling in soma (**pentagons**), proximal (**black arrowheads**), and distal dendritic fields (**white arrowheads**). IHC provided exceptional visualization of labeling in proximal dendritic fields. Scale bar in **b** (5 mm) applies to **a** and **i-j**; scale bar in **c** (200 μm with 5× objective) applies to **d-e** and **k-m**; scale bar in **f** (50 μm at 20×) applies to **g-h** and **n-p**. GFP and RFP were visualized with fluorescence microscopy for Epi and IF or bright field microscopy for IHC. Z-stacks: e = 20z; f = 20z; g = 23z; l = 18z; m = 15z; n = 17z.

**Figure 7.**
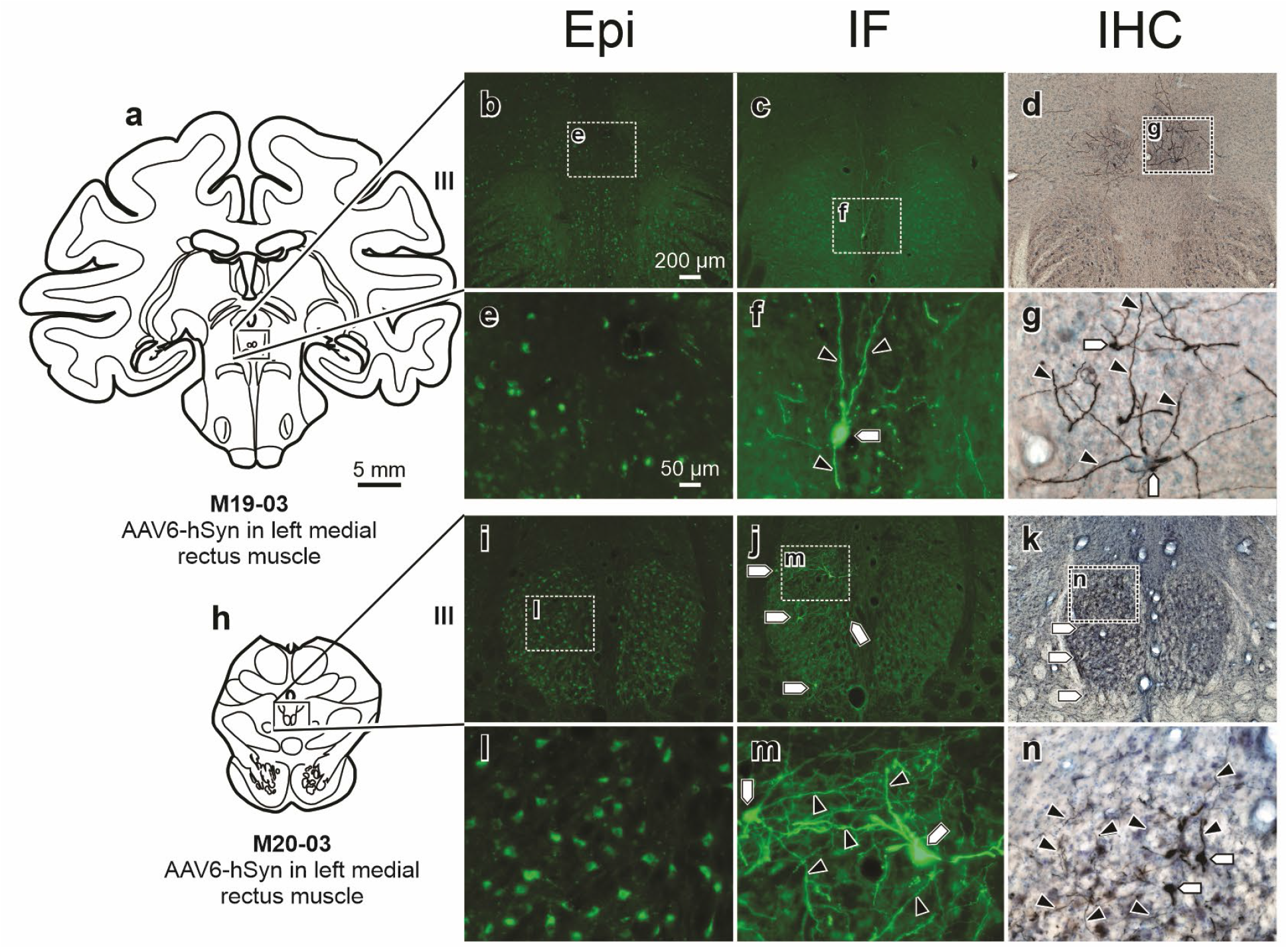
Negligible fluorescent expression was detected in motoneurons using Epi, but robust labeling was observed using IF and IHC following eye muscle injections. (**a**) Low magnification drawing of III following injection of AAV6-hSyn-ChR2(H134R)-eYFP in left medial rectus muscle of M19-03. Epi (**b, e**), IF (**c, f**), and IHC (**d, g**) sections for M19-03 III. (**h**) Low magnification drawing of left III following injection of AAV6-hSyn-ChR2(H134R)-eYFP in left medial rectus muscle of M20-03. Epi (**i, l**), IF (**j, m**), and IHC (**k, n**) sections for M20-03 III. Labeling of dendritic fields (**arrowheads**) and soma (**pentagons**) was observed only when tissue was assessed using IF or IHC. Scale bar in **a** (5 mm) applies to **h**; scale bar in **b** (200 μm at 5×) applies to **c-d** and **i-k**; scale bar in **e** (50 μm at 20×) applies to **f-g** and **l-n**. GFP was visualized with fluorescence microscopy for Epi and IF or bright field microscopy for IHC. Z-stacks: e = 20z; f = 23z; g = 13z; l = 20z; m = 20z; n= 17z.

In addition to V2, we also examined visual areas MT and MST (MT/MST) (**Fig. 5i-o**) from the same SC injection. As was observed in V2, Epi (**Fig. 5j, m**) provided a moderate capacity to detect retrogradely labeled cell bodies (**Fig. 5m; pentagons**) and apical dendrites (**Fig. 5m; black arrowheads**) in MT/MST. Both IF (**Fig. 5k, n**) and IHC (**Fig. 5l, o**) resulted in enhanced ability to visualize soma (**Fig. 5n, o; pentagons**), along with dense proximal (**Fig. 5n, o; black arrowheads**) and distal (**Fig. 5k, l; white arrowheads**) dendritic field labeling. To determine if this was just a function of neurons with projections to the SC, we examined a second case in which rAAV2-retro-CAG-tdTomato was injected into the left FEF (**Fig. 5p**). This injection retrogradely labeled neurons in MT/MST having projections to the FEF (**Fig. 5q-w**). Epi (**Fig. 5r, u**) revealed sparse labeling in MT/MST neuronal soma (**Fig. 5u; pentagons**) and proximal (**Fig. 5u; black arrowheads**) dendritic fields. In contrast, IF (**Fig. 5s, v**) and IHC (**Fig. 5t, w**) revealed enhanced labeling of soma (**Fig. 5v, w; pentagons**) and proximal dendritic fields (**Fig. 5v, w; black arrowheads**) of the MT/MST neurons.

The most frequent observation was that labeling in projecting neuronal populations was negligible using Epi assessment, but present using IF or IHC methods of assessment (**Fig. 6 and 7**). For example, following injection of rAAV2-retro-CAG-tdTomato into the right SC (**Fig. 6a**, same injection as illustrated in Figure 5a), we assessed retrograde labeling in the right lateral intraparietal cortex (LIP) (**Fig. 6b-h**). Epi assessment (**Fig. 6c, f**) revealed no detectable labeling, whereas IF (**Fig. 6d, g**) and IHC (**Fig. 6e, h**) assessments revealed virally mediated labelling in cell bodies (**Fig. 6g, h; pentagons**), basal and proximal dendrites (**Fig. 6g, h; black arrowheads**), and distal dendritic fields (**Fig. 6d, e; white arrowheads**). In another case in which rAAV2-retro-CAG-GFP was injected into right SC (**Fig. 6i**), we assessed expression in the right FEF (**Fig. 6j-p**). Epi (**Fig. 6k, n**) was insufficient to visualize retrogradely labeled corticotectal neurons in FEF. However, following amplification with IF (**Fig. 6l, o**) or IHC (**Fig. 6m, p**), transgene expression was detected in cell bodies (**Fig. 6o, p; pentagons**), basal and proximal dendrites (**Fig. 6o, p; black arrowheads**), and distal dendritic fields (**Fig. 6l, m; white arrowheads**). In both cases, IHC produced particularly robust labeling of proximal dendritic fields (**Fig. 6h, p; black arrowheads**).

Next, we evaluated expression in retrogradely transduced motoneuronal populations of the third cranial nerve, the oculomotor nucleus (III), following injections of AAV6 into extraocular muscles. The overwhelming majority of eye muscle injections resulted in negligible labeling using Epi. The fluorescence that was observed in the Epi photomicrographs (**Fig. 7b, e, i, l**) was background. Following injection of AAV6-hSyn-hChR2(H134R)-EYFP in the left medial rectus, Epi assessment (**Fig. 7b, e**) of the left III revealed no detectable labeling, while IF (**Fig. 7c, f**) and IHC (**Fig. 7d, g**) provided robust visualization of motoneuronal cell bodies (**Fig. 7f, g; pentagons**) and dendritic fields (**Fig. 7f, g; black arrowheads**). The same results were found after injection of AAV6-hSyn-hChR2(H134R)-EYFP into the left medial rectus of a second macaque (**Fig. 7h-n**).

In summary, the data from CNS and PNS injections demonstrated that Epi assessment produced highly variable results when used for visualization of viral mediated transgene expression, whether locally, anterograde, or retrograde to the injection. Conversely, immunostained tissue provided more reliable and comprehensive visualization of neuronal labeling, regardless of the viral construct or conditions used.

As a final step in comparing the reliability and validity of Epi vs. immunostaining for visualizing virally mediated transduction in the primate CNS, we performed opposite color amplification to visualize red Epi and green IF within the *same* sections. Following an injection of rAAV2-retro-CAG-tdTomato in the right SC (**Fig. 8a**), viral mediated labeling was evaluated in the right nucleus of the brachium of inferior colliculus (BIC) (**Fig. 8b-h**), the central mesencephalic reticular formation (cMRF) (**Fig. 8b, i-n**), and the FEF (**Fig. 8o-u**). Within BIC, labeled axonal tracts were clearly visible upon Epi assessment (**Fig. 8c, e, f, h**) and following IF (**Fig. 8d, e, g, h**). In this example, the axonal labeling under Epi was particularly intense. Following IF amplification, the intensely labeled axons observed in Epi were not clearly visible, but a plethora of finely labeled neuronal processes was visible. This dichotomy could be attributed to two things. First, there could be weak antibody penetrance through the 75 μm thick tissue. Second, available antibody could be quenched near the surface of the section. This means that most available antibody would bind to tdTomato in the fine neuronal processes near the surface of the section, leaving little antibody to bind to antigen deep within the tissue section. In the cMRF, Epi (**Fig. 8i, k, l, n**) revealed scant labeling in dendritic fields (**Fig. 8l, n; white arrowhead**), whereas IF (**Fig. 8j, k, m, n**) provided robust labeling in somas (**Fig. 8m, n; black pentagons**) and dendritic fields (**Fig. 8m, n; black arrowheads**). Finally, Epi assessment (**Fig. 8p, r, s, u**) of the retrogradely labeled corticotectal projecting FEF neurons revealed minimal labeling in cell bodies (**Fig. 8s, u; white pentagons**) and apical dendrites (**Fig. 8s, u; white arrowheads**). Conversely, IF (**Fig. 8q, r, t, u**) provided robust visualization of somas (**Fig. 8q, t, u; black pentagons**), proximal dendritic fields (**Fig. 8t, u; black arrowheads**), and distal dendritic stem (**Fig. 8q; black arrows**). Importantly, amplification revealed more transduced neurons in both cMRF and FEF and more neuronal features for individual neurons in all three regions (fine axonal labeling in the BIC and dendritic labeling within the cMRF and FEF).

**Figure 8.**
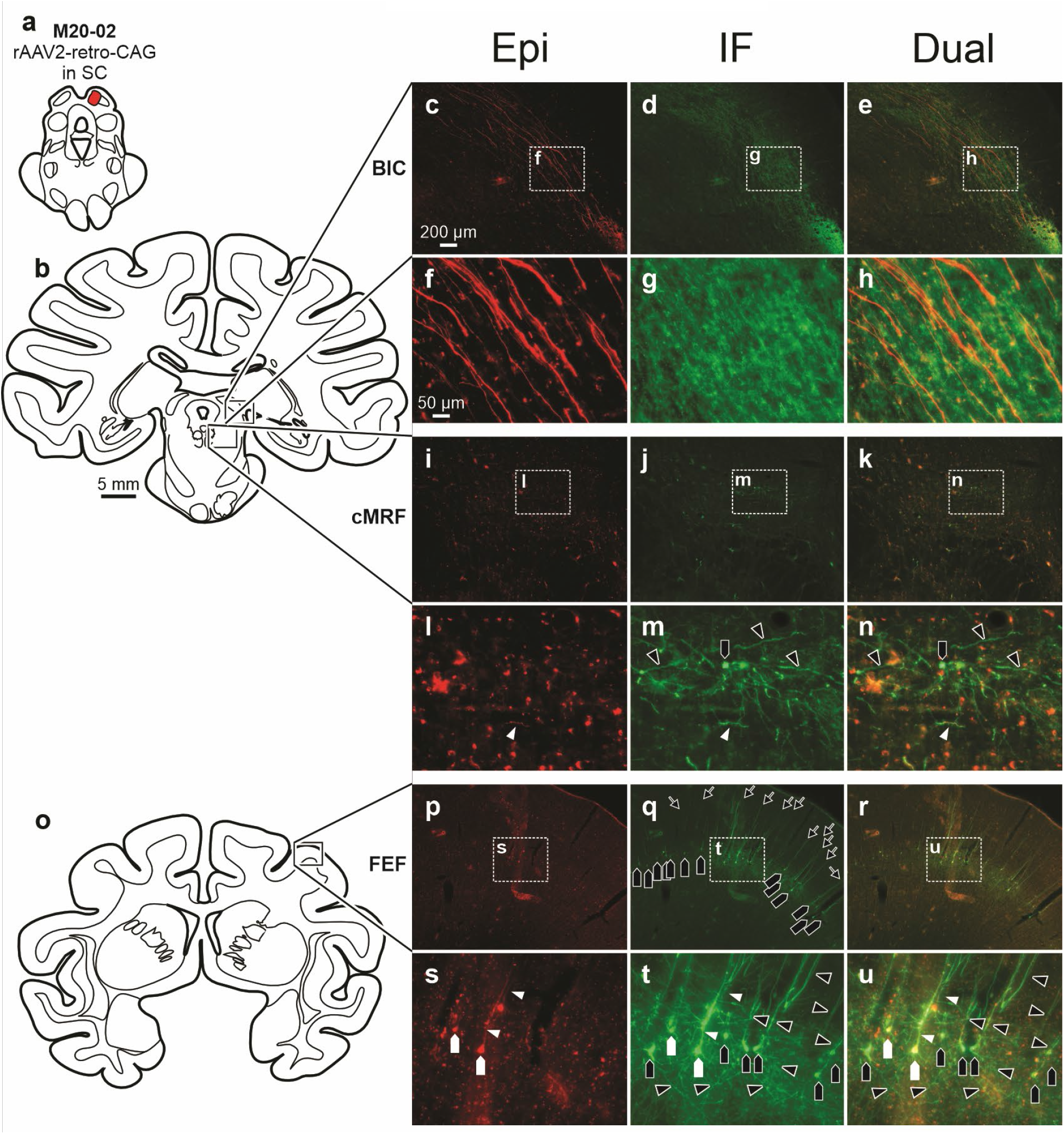
Direct comparison of Epi and IF demonstrated that labeling was more consistent and robust with amplification than without amplification. **(a)** Low magnification drawing illustrating the injection of rAAV2-retro-CAG-tdTomato in right SC of M20-02. This injection resulted in detectable labeling within the BIC (**b, c-h**), cMRF (**b, i-n**), and FEF (**o, p-u**). Epi labeling is red, while IF is green. (**b**) Low magnification drawing of the section illustrating the approximate location of the right BIC (top box) and cMRF (bottom box) from which the photomicrographs were taken. Within the BIC, labeled axons were clearly visible in Epi (**c, f**), while finely labeled neuronal processes were detectable following IF amplification (**d, g**). The merged images show dual labeling (**e, h**). Within the cMRF, scant labeling was present using Epi assessment (**i, l**) with the exception of some neuronal structures, for example, a small portion of a labeled dendrite (**white arrowhead**). Conversely, using amplification (**j, m**), robust labeling of soma (**black pentagons**) and dendritic fields (**black arrowheads**) was observed. (**o**) Low magnification drawing illustrating the location of photomicrographs in the right FEF of M20-02. Epi (**p, s**), IF (**q, t**), and Dual (**r, u**). Again, in the assessment of corticotectal neuronal labeling within FEF, minimal labeling was observed in soma (**white pentagons**) and apical dendrites (**white arrowheads**) using Epi assessment, whereas dense labeling of soma (**black pentagons**), basal and apical dendrites (**black arrowheads**), and distal dendritic fields and tufts (**black arrows**) were revealed using IF. Scale bar in **b** (5 mm) applies to **a** and **o**; scale bar in **c** (200 μm at 5×) applies to **d-e, i-k**, and **p-r**; scale bar in **f** (50 μm at 20×) applies to **g-h, l-n,** and **s-u**. RFP and GFP were visualized with fluorescence microscopy. Z-stacks: e = 25z; f = 25z; g = 25z; k = 15z; l = 15z; m = 15z; r = 20z; s = 20z; t = 20z.

This study focuses on the use of AAV2 derivatives for central injections and AAV6 for peripheral injections. Although beyond the scope of this study, additional experiments demonstrated that the observed differences between Epi and immunostaining extend to other viral serotypes, promoters, and fluorophores. In pilot experiments, tissues from additional animals were processed using only IHC to compare with Epi. These additional cases included a total of 28 injections (19 craniofacial intramuscular injections and 9 CNS injections) in 10 additional macaques using the following viral vectors: AAV1-Syn-ChrimsonR-tdTomato, AAV1-hSyn-HI-eGFP-Cre-WPRE-SV40, AAV2-EF1α-mCherry-IRES-WGA-Cre, rAAV2-retro-CAG-GFP, rAAV2-retro-CAG-tdTomato, AAV5-EF1a-eGFP-WPRE-rBG, AAV5-EFlα-Ace-mNeon-4AA, AAV6-hSyn-hChR2(H134R)-EYFP, AAV9-CAG-ChR2-GFP, AAV9-hSyn-iLMO-EYFP, and AAV10-hSYN-nucGFP-2A-LOX-nucRFP-LOX. Since the current report compares Epi to both methods of immuno-amplification and primarily focuses on AAV serotypes 2 and 6, these pilot cases were excluded. Critically, these data revealed the same generalized patterns reported above for Epi versus immunostaining (not shown).

## DISCUSSION and conclusion

In the present study, we compared Epi and immunostaining techniques (IF and IHC) for their ability to detect AAV transduction in several tissues. Anatomical analysis of AAV constructs included regions local to the injection site and regions known to have projections to or from the injected sites. For each population, we compared the abilities of different histological approaches to detect and distinguish neuronal structures (i.e., somas, dendritic fields, axons, and axon terminals) labeled via viral transduction. In the best-case scenarios, Epi assessment revealed moderate transgene expression that was occasionally present local to the central injection sites (**Fig. 1, 2**). More frequently, however, Epi showed negligible labeling at intraparenchymal injection sites (**Fig. 1, 3**) and in the axon terminal fields efferent from the injection sites (**Fig. 4**). In a few instances, after retrograde viral labeling, some cells were detected with Epi (**Fig. 5**), but more often the detection of retrogradely labeled neurons was negligible (**Fig. 6-8**). When Epi identified labeled cells, the fluorescence was detectable only within the soma, proximal, and apical dendrites.

In contrast, immunostaining reliably produced enhanced visualization of transgene expression at intraparenchymal injection sites (**Fig. 2 and 3**), in anterogradely labeled terminal fields (**Fig. 4**), and in retrogradely labeled neuronal populations (**Fig. 5-7**). In IF and IHC amplified tissues, antibody labeling against the expressed exogenous protein was observed within most neuronal compartments, including the soma, axons, terminal fields, and throughout the dendrites. Importantly, IF and IHC amplification consistently provided visualization of expression in regions where Epi assessment provided minimal detection of expression. Similar results were observed in cranial nerve nuclei following AAV injections in the extraocular muscles (**Fig. 7**). Finally, a direct comparison of Epi to IF amplification in the same tissue revealed that differences in detectable viral transgene expression were the result of amplification, rather than variance between sections (**Fig. 8**).

The implications of these observations are of particular concern for studies that depend on viral vectors to deliver exogenous genes to targeted neuronal populations in primates and potentially reflect similar limitations for clinical gene therapies.

### Technical considerations

In the present study, IF and IHC methods were used to amplify the fluorescence conferred by the viral genome. Thus, it was unsurprising that visualization of transgene expression was enhanced following IF and IHC. However, the degree to which amplification provided enhanced visualization of both the number of neurons and the morphological details within individual neurons was striking, especially in comparing Epi and IF results within the same tissue (**Fig. 8**). Furthermore, the variety of conditions used demonstrated that differences between Epi and IF were unlikely produced by factors such as survival time, injection volume, titer, multiplicity of infection, viral capsid, promoter, or transgene. It is possible that Epi and immunostaining each provide unique insights into the efficacy of viral transduction and gene expression.

#### Epifluorescence microscopy

Epi is the simplest approach since all that is required is to section the brain and mount the tissue to slides for microscopic analysis. A major benefit of Epi is that it provides a raw metric for the degree of transgene expression within transduced neurons. The major downside of Epi is that it lacks sensitivity for assessing the extent of viral transduction (i.e., number of infected neurons) and localization of expression (i.e., somas versus dendrites). Relying solely on Epi assessment could therefore lead to false-negative conclusions about whether a viral construct succeeded in transduction of a given brain region.

#### Immunofluorescence and immunohistochemistry

The main benefits of IF and IHC include visualization of a greater number of transduced cells, assessment of projection labeling, and enhancement of morphological features for single labeled cells. Specifically, IF and IHC are frequently required to visualize small, discrete components of neurons including the complete dendritic arbors or terminal fields. Compared with Epi, the main downsides of IF and IHC are that they are more technically demanding and have added costs for reagents and personnel effort. In comparing immunostaining techniques, IF is optimized for multiplexing, which is more difficult with IHC since there are fewer easily distinguishable colors available. However, labeling produced using IHC has a longer shelf-life compared with fluorescence from IF, which is more susceptible to bleaching over time ^24^.

In summary, the present study demonstrates that the best way to gain insight into viral gene expression is to use both Epi and immunostaining techniques or to multiplex within single sections as illustrated in **Figure 8**. Because of the differences between each anatomical method for assessing viral expression, the choice of approach will largely depend upon one’s experimental goals. Regardless, anatomical assessment is increasingly important as newly optimized features arise from capsid engineering and genomic modifications and as our understanding of basic virology evolves ^17,18,25–31^. Generally, it can be concluded that processing tissues using multiple approaches maximizes the information gleaned from any individual sample. This is particularly important in the primate model, for which there is little, systematic information about viral vector efficacy at present.

The results from this study demonstrate that an amplification step is currently necessary to visualize the extent of virally mediated transgene expression in macaques. Just because a gene product is detectable by immunostaining, however, does not mean that it has accumulated to the concentration required for manipulation using neuronal actuators. Hence, a potential downside of immunostaining may be in generating the false impression that expression is substantial when it is in fact too low to influence neuronal activity. It is possible that transgene labeling visualized using Epi better reflects the amount of expression necessary for physiological or behavioral efficacy of neuronal actuators. Thus, we hypothesize that Epi provides insight to the efficacy of neural actuators that are included in the viral construct, while IF and IHC can amplify even minimal transduction to yield comprehensive information about gene expression across neural compartments. Additional physiological and behavioral studies are necessary to understand how anatomical assessment of transgene expression reflects physiological or behavioral efficacy of neuronal actuator proteins.

### Potential mechanisms

The present study was retrospective, so investigating the mechanisms that underlie the observations is beyond its scope. However, the observations herein raise questions about mechanisms underlying the lackluster transgene expression observed in the primate brain. Grossly, poor expression in the primate model (compared to cell culture and rodents) may arise from three categories: 1) differences in transduction, 2) immunological responses (adaptive and innate), and 3) model validity.

#### Transduction and immunological responses

Studies in cell culture and rodent models have revealed that loss of capsids throughout transduction and immunological neutralization are established barriers ^32^. Compounding the problem, little has been done to make direct comparisons of transduction across models (i.e., cell culture vs. rodent vs. primate). Given these facts, primates may be more efficient at reducing the number of infective viral units along the transduction pathway. In addition, there is considerable divergence between the immune systems of different species ^33^, which differentially impacts immunological responses to viral or bacterial exposure ^34^. As viral methods are further refined for primates, variables such as codon optimization will become increasingly critical for ensuring strong expression through a minimization of immunogenic features in the viral genome and transgenic protein ^35^.

#### Model validity

Fundamentally, it is necessary that the bioengineering of viral vectors has the required validity. While more cost effective, will engineering and optimizing viral constructs in phylogenetically lower species translate to primates? The present results suggest that current constructs lack face validity (i.e., capsid efficacy fails to translate from cell culture/rodents to primates) and construct validity (i.e., the capsid functions differently in primates than it does in other species) ^31,36–39^.

### Comparison with prior studies

How do the present results in macaque compare to similar studies done in rodents? A recent report in mice found that, given a 20 to 29-day expression period following hypothalamic AAV delivery of genes encoding mCherry, IHC amplification was required to visualize the fluorescent transgene ^40^. Another study examined the time course of transgene expression following striatal injections of AAV1, 2, 5, and 8 in rats ^41^. The results revealed that GFP could be detected as early as 4 days with IHC amplification, but fluorescence was undetectable by Epi before 7 days. By 4 weeks, the Epi-detected fluorescent signal was comparable to IHC detection ^41,42^. The present study builds on previous findings using several recombinant AAV parental and chimeric serotypes and survival durations ranging from 51 to 147 days post-injection. However, unlike results from Reimsnider et al. (2007), results from primates that are presented here suggest that Epi and immunostained tissue provide different impressions of the efficacy of viral mediated transgene expression, even following long survival durations. This raises the possibility that, when using the current generation of viral constructs, survival durations beyond 147 days may be required in primates to achieve comparable detection of transgene expression for Epi and immunostaining assessments.

### Potential implications for differences in gene expression across species

In addition to differences in anatomical assessment of viral technologies across species, there are discrepancies in the functional efficacy of neuronal actuators and indicators between rodent and primate models. This divergence is best exemplified by the rapid adoption and implementation of effective neuronal actuator and indicator techniques in rodents, where such approaches have been successfully applied to most domains of neuroscientific research. On the other hand, these techniques have not been widely implemented in non-human primate models. The primate neuroscientific community has only scantly published on neuronal actuator and indicator techniques ^1–3,43,44^. Furthermore, actuator and indicator efficacy remains largely unreliable and ineffective in one or more of the important categories for defining its success (i.e. anatomical, physiological, and behavioral) ^2,4^.

The differences in efficacy between primates and rodents could be due to several factors. The simple explanation is that widespread implementation of viral technologies in primates is hindered due to long gestation periods, small litter sizes, high costs, and ethical roadblocks. These limitations are reduced in rodent species. The results from the current study suggest an alternative explanation for the lack of actuator efficacy in the primate model. If transduced cells have not accumulated enough exogenous transgene, as reflected by minimally detectable fluorescence visualized using Epi assessment, there may not be sufficient mature actuator proteins present to modify a given cell’s membrane potential. This may explain why, despite being anatomically detectable with immuno-amplification, macaque neurons fail to respond in the presence of the appropriate trigger (i.e., proper wavelength of light in the case of opsins or ligand in the case of chemogenetic receptors) ^2^. A general failure to modify neuronal activity would then lead to a failure to modify both circuit activity and behavior. These results suggest that the field faces a challenge that must be addressed. Namely, neuronal transduction and exogenous gene expression driven by viral vectors need to be enhanced in primates.

The present work is not the first to suggest species differences in AAV-driven transgene expression. In an elegant and carefully performed series of experiments, ^45^ compared the location of different DREADDs (Designer Receptors Exclusively Activated by Designer Drugs) conjugated with mCherry in mice and monkeys. The authors demonstrated that an hM4Di-mCherry construct was successfully transported to the plasma membrane of neurons in mice, but not in primates. The hM3Dq-mCherry construct, in contrast, was largely confined to the plasma membrane of monkey neurons. Thus, gene expression and protein localization depended on both the viral construct and animal model. In another study, Chen et al. (2022) evolved AAV capsids in mice to selectively target dorsal root ganglion. In rodents, the capsids targeted ganglion cells and not the CNS. In contrast, in two primate species, the capsids crossed the blood brain barrier and transduced neurons within the CNS ^31^. Such observations suggest a distinction in how cells of different species interact with viral vectors to express exogenous transgenes. These findings highlight the need to continue to develop and optimize viral constructs, neuronal actuators, and indicators *within* the primate model.

This also highlights the need to understand the mechanisms underlying such species differences. Primate and rodent immune systems are different ^33^. Thus, potential toxicity from the capsid, viral genome, and transgene products are quickly recognized and eliminated by neutralizing antibodies in non-human primates (unpublished observations; ^46,47^) or through intracellular immune responses ^32^. Additionally, there may be subtle differences in transcription or translation of the viral genome between species, which ultimately lead to distinct transgene expression patterns ^17,45^. Further studies will be required to understand how species differences impact viral transduction and gene expression.

### Relevance to gene therapy

The potential species differences emphasize the importance and utility of the primate model in preclinical gene therapy. The ability to deliver an exogenous gene to a target cellular population and then reliably drive transgene expression in primates is clinically relevant. The present study establishes that viral-mediated expression in primate brain tissue is low as assessed using Epi. The necessity for amplification of the reporter suggests a need for improved viral constructs that will provide enhanced transgene expression in the primate model. Further, this suggests that the current generation of viral constructs is potentially not optimized for clinical use in humans.

Significant advances have been made to improve the efficacy of viral vectors and neuronal actuators and indicators, but their development and *in vivo* optimization has predominantly been performed in rodent or non-mammalian species ^22,27,48^. This is the conventional pharmaceutical pipeline for developing treatments, but for viral vectors it has its drawbacks. A recent example was the overly evolved AAV-PHP.B for the C57BL/6J mouse strain ^39^ which failed to translate to primates and even to other mouse strains ^36–38^. It is therefore possible that there is over-optimization of reagents that are biasing successful outcomes toward rodents. Thus, both the basic sciences and clinical therapies will benefit from developing and accurately assessing viral delivery approaches in the primate, for the primate.

## Acknowledgments

We thank Jessi Cruger and the veterinarians and staff of Duke’s Division of Laboratory Animal Resources for animal care and surgical technical assistance. We thank Dr. Greg Field for valuable feedback on early versions of this manuscript.

## Contributions

T.B.D., H.G.E., M.A.B., E.J.J., A.R.B., M.A.S., and M.O.B. conducted experiments. T.B.D. and M.O.B. wrote the manuscript. M.A.S. and R.J.S. edited the manuscript.

## ABBREVIATIONS

III: Oculomotor nucleus
BIC: Brachium of the inferior colliculus
cMRF: Central mesencephalic reticular formation
FEF: Frontal eye field (Area 8)
ia: Inferior arcuate sulcus
ips: Intraparietal sulcus
LIP: Lateral intraparietal cortex
MD: Mediodorsal thalamic nucleus
MIP: Medial intraparietal area
MST: Medial superior temporal area
MT: Area MT (V5)
PE: Parietal area PE
sa: Superior arcuate sulcus
SC: Superior colliculus
V2: Secondary visual cortex

